# On the Mathematics of Populations

**DOI:** 10.1101/612366

**Authors:** M. Gabriela M. Gomes

## Abstract

Selection acting on unobserved heterogeneity is a fundamental issue in the mathematics of populations. As recognised in disciplines as diverse as demography [11, 23, 24], ecology [10, 9], evolution [21] and epidemiology [1, 4, 5], in any population, individuals differ in many characteristics and it is essential that researchers understand which of these are under selection and how selection processes operate. Here I describe conceptual and methodological developments in demography and ecology, and discuss the importance of adopting similar approaches in evolution and epidemiology.

## 1 Demography

Unobserved heterogeneity can result from many interacting genetic and environmental factors. When operating on individual survival, it modifies the composition of cohorts as they age [11, 23]. Frail individuals die younger, leaving the cohort progressively composed of those who are more robust and have a propensity to live longer. This form of selection acting on longevity operates within cohorts and distorts patterns of age-specific survival [24]. It may also affect other life-history traits via correlations with longevity. For example, if those individuals who live longer have lower fecundity, then selection operating on individual longevity will reduce fecundity at the population level.

## 2 Ecology

Demographic heterogeneity has been contrasted with demographic stochasticity in ecology [10, 9]. Demographic heterogeneity, defined as variation among individuals in life-history characteristics, has been addressed in ecology by structured population models [3, 19]. The approach consists of incorporating the most important differences into the individual state of a matrix model. A major challenge is to reconcile those individual differences that matter (given the question under study) with those that can actually be measured. A natural tendency is to account for characteristics that can be measured most easily, such as age, size, or major developmental states, and collapse other important sources such as genetic variation, spatial heterogeneity in the environment, and differential exposure to stressors, under some form of unmeasured stochasticity. Recent research has elucidated, however, that demographic heterogeneity and demographic stochasticity have opposing effects in population dynamics and cannot be modelled interchangeably [3, 19].

## 3 Evolution

Recent research has begun to emphasise the importance of considering individual variation in non-heritable fitness components when interpreting the results of evolutionary studies [21]. By accommodating explicitly for individual variation in non-heritable fitness components, we have shown how common proxies for genotype fitness may be affected by a form of selection that is invisible to evolution and how this may explain observed trends when fitness is measured across stress gradients [6]. We then propose that unaccounted phenotypic variation within genotypes is capable of stabilising coexistence of multiple lineages and unexpectedly affect patterns of genetic variation, especially when levels of stress fluctuate.

Understanding the selective forces that shape variation is at the heart of evolutionary theory. Selection acting on non-heritable characteristics has not been given full attention, either because it is not directly measurable or because it was believed to be inconsequential for evolution. In [6] we oppose these commodities and argue that selection on non-heritable fitness components is essential both as a natural step in theory development and as a necessity for accurate interpretation of population and quantitative genetic data. More practically, we propose concrete experimental designs across stress gradients for informing models of variation and selection.

## 4 Epidemiology

Variation in individual characteristics has a generally recognised impact on patterns of occurrences in populations, and occurrence of disease is no exception. In infectious diseases, the focus has been on heterogeneities in transmission through their nonlinear effects on the basic reproduction number, *R*_0_, in ways which are unique to these systems [15, 25]. The need to account for heterogeneity in disease risk, however, is not unfamiliar in epidemiology at large, where frailty terms are more generally included in linear models to improve data interpretation [1].

The premise is that variation in the risk of disease (whether infectious or not) goes beyond what is captured by measured factors, and a distribution of unobserved heterogeneity can be inferred from incidence patterns in a holistic manner. Such distributions are needed for eliminating biases in interpretation and prediction, and can be utilised in conjunction with more common reduction-ist approaches, which are required when there is desire to target interventions at individuals with specific characteristics.

To contrast different forms of heterogeneity, we consider four versions of a *Susceptible-Infected* (*SI*) model in a population of constant size.

### Homogeneous *SI* model

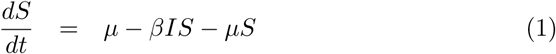

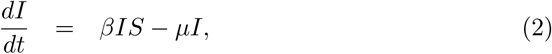

where *S* and *I* are the proportions of susceptible (uninfected) and infected (infectious) individuals, respectively, *β* is a transmission coefficient (effective contact rate) and *µ* accounts for birth and death. The force of infection upon uninfected individuals is *λ* = *βI* and the basic reproduction number is:

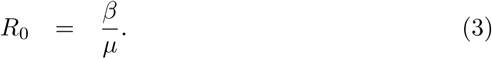

### *SI* model with heterogeneous susceptibility

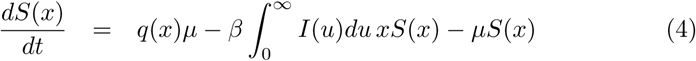

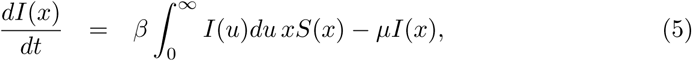

where *x* is an individual susceptibility factor taking values on a continuum, *q*(*x*) is a probability density function, *S*(*x*) and *I*(*x*) represent the densities of susceptible and infected individuals, respectively, and *β* and *µ* are parameters governing transmission and demographic processes as before. The force of infection upon the average uninfected individual is 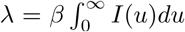, which is then affected by a factor *x* to conform with individual susceptibilities. The basic reproduction number for this system is:

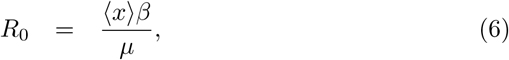

where ⟨*x*⟩ is the first moment of the susceptibility distribution (or mean susceptibility, i.e. 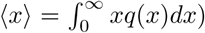. All model solutions presented here assume ⟨*x*⟩ = 1, in which case the *R*_0_ expression of the heterogeneous susceptibility model coincides with the homogeneous.

Figure 1 shows the prevalence of infection over time as governed by models (1)-(2) and (4)-(5), as well as density plots (frequency in the homogeneous case) for the susceptible and infected compartments at three time points: before the start of the epidemic; at the endemic equilibrium corresponding to 20% prevalence; after 100 years of control. The control measure simulated in Figure 1B is the 90-90-90 treatment as prevention cascade advocated by UNAIDS for HIV, which stipulates that 90% of infected individuals are detected, 90% of those detected enter antiretroviral therapy, and 90% of those entering treatment achieve viral suppression, becoming effectively non-infectious. As a result of this cascade, transmission is reduced by approximately 73%.

**Figure 1:**
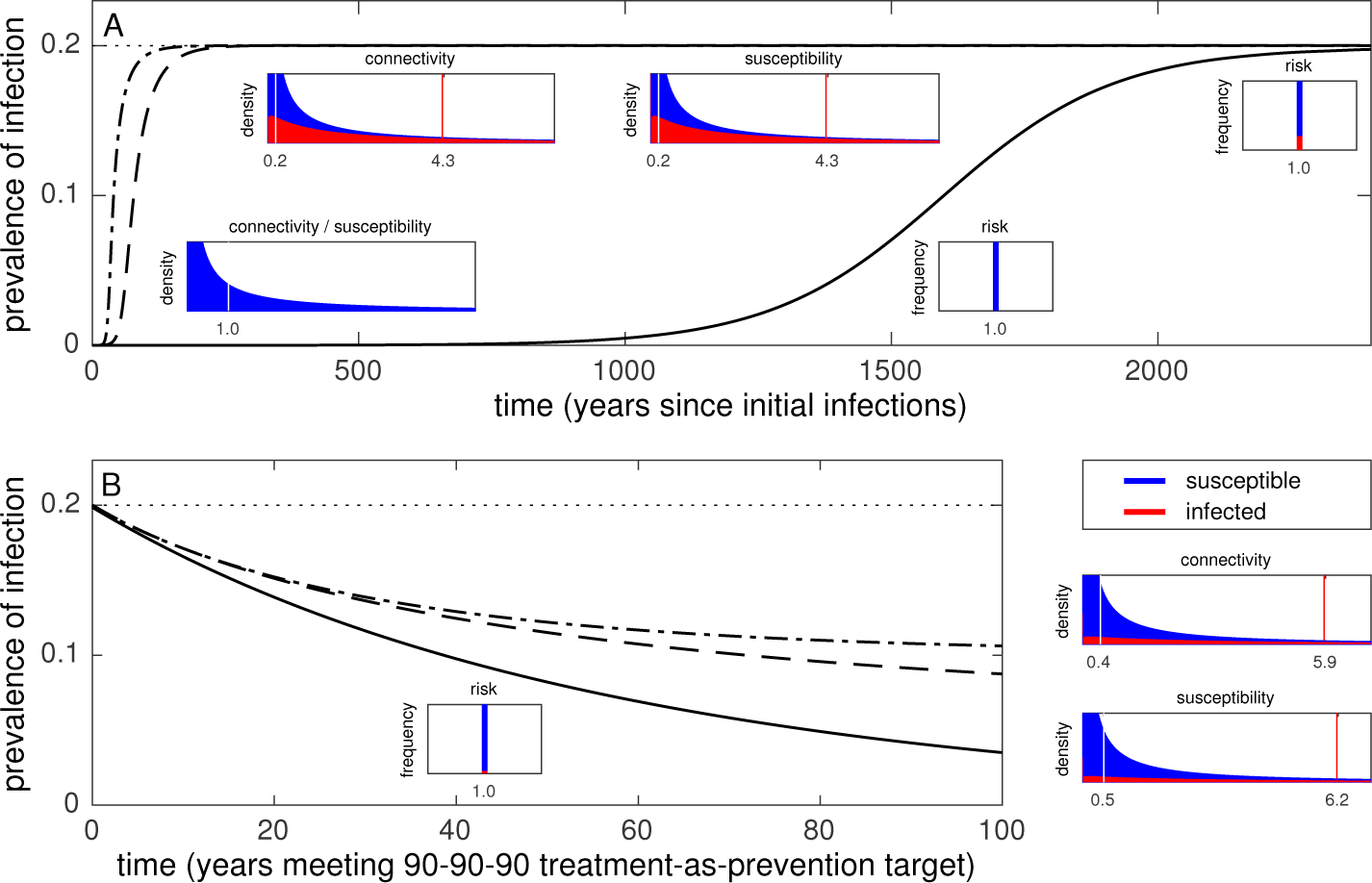
Prevalence trajectories under models (1)-(2) and (4)-(5). Two susceptibility distributions are simulated: homogeneous (solid black trajectories); gamma distributed susceptibility to infection (dashed black trajectories). Insets depict the gamma distribution with mean 1 and variance 10 utilised in the heterogeneous implementation side-by-side with its variance 0 counterpart implicit in the homogeneous model. The vertical white and red dotted lines added to the gamma distribution plots mark mean values for the susceptible (blue) and infected (red) subpopulations, respectivelly.

In comparison with the homogeneous model, a higher *R*_0_ is required to reach the same endemic level, and the same control measure has lower impact under heterogeneity (this is irrespective of the *R*_0_ expressions (3) and (6) being the same). This is evidenced by comparison of the solid (homogeneous) and dashed (heterogeneous) trajectories, and explained by the density plots. As the infection spreads in the population, more susceptible individuals are likely to be infected earlier (red distributions, with higher mean - red dotted lines) and, consequently, those who remain uninfected are being selected for lower susceptibility (blue distributions, with lower mean - white dotted lines). This process slows down the epidemic since the mean susceptibility among those at risk effectively decreases as the epidemic progresses. As a result, the heterogeneous model requires higher values of *R*_0_ to attain the same endemic level as its homogeneous counterpart, and becomes more resilient to interventions designed to reduce transmission. Figure 1 was generated assuming a gamma distribution with mean 1 and variance 10 for the heterogeneous susceptibility model, but the effects described above are generally manifested with a strength that increases with the variance (or coefficient of variation).

### *SI* model with heterogeneous infectiousness

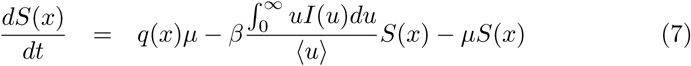

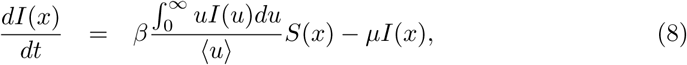

where *x* is an individual infectiousness factor taking values on a continuum, and *q*(*x*), *S*(*x*), *I*(*x*), *β* and *µ* represent densities and parameters as before. The force of infection upon uninfected individuals is 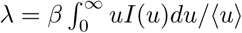, and the basic reproduction number is the same as in the homogeneous implementation.

### *SI* model with heterogeneous connectedess

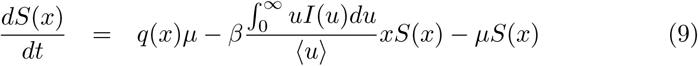

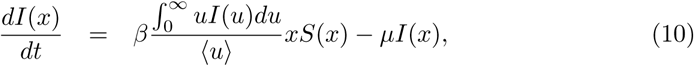

where *x* is again a factor taking values on a continuum but now affecting both dispositions for acquiring infection and for infecting others, due to variable connectedess. Everything else is defined as before, and the basic reproduction number is:

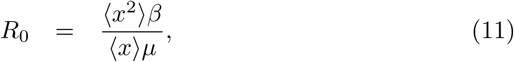

where ⟨*x*^2^ ⟩ is the second moment of the connectedess distribution (i.e. 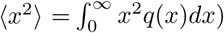. The expression for *R*_0_ arising from this model is typically different from the other implementations, even when ⟨*x* ⟩= 1.

Figure 2 shows the prevalence of infection over time as governed by models (7)-(8) and (9)-(10), accompanied by density plots for the susceptible and infected compartments before the start of the epidemic, at the 20% prevalence equilibrium, and after 100 years of control as above.

**Figure 2:**
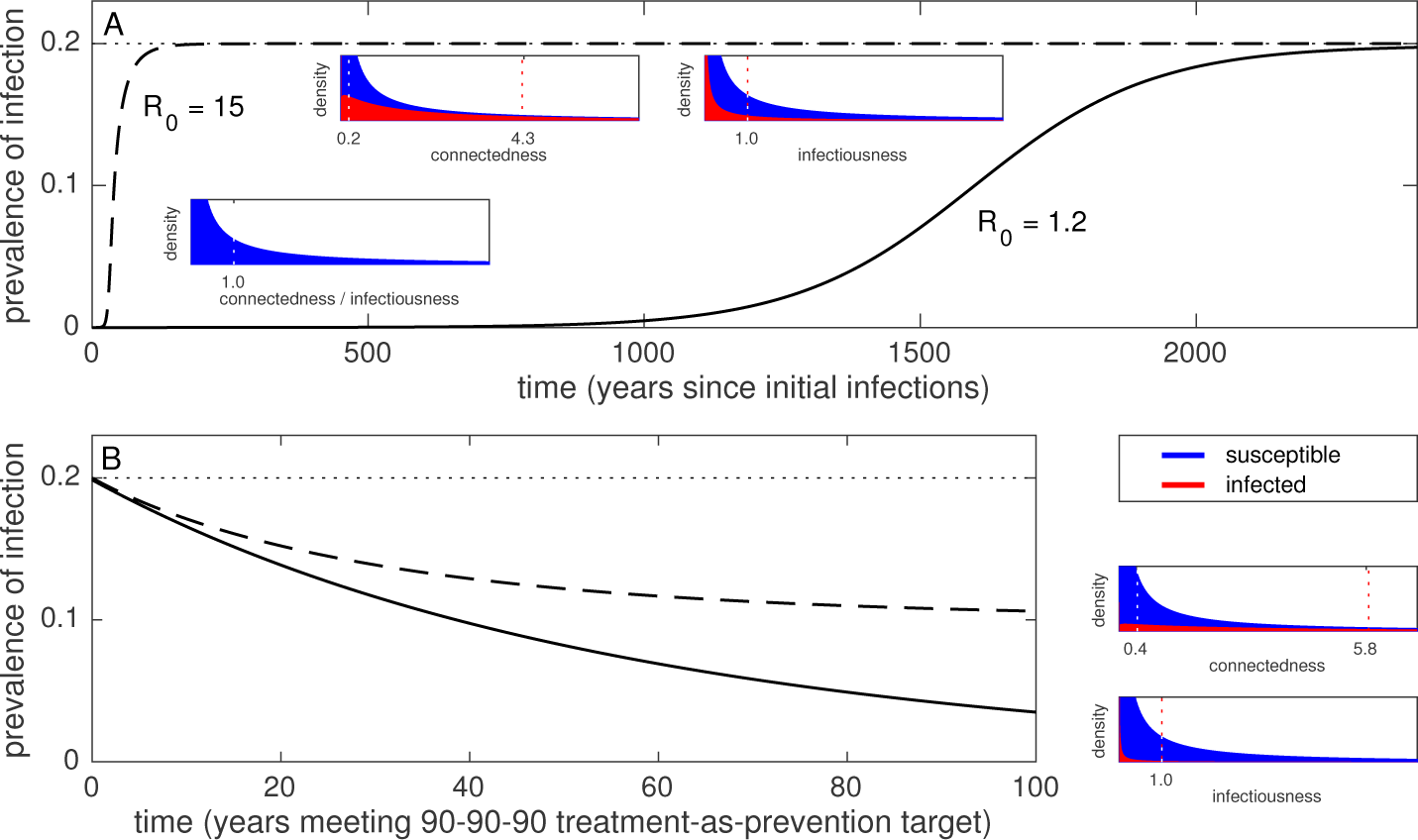
Prevalence trajectories under models (7)-(8) and (9)-(10). Two distribution implementations are simulated: gamma distributed infectiousness (solid black trajectories); gamma distributed connectedess (dashed black trajectories). Insets depict the gamma distribution with mean 1 and variance 10 utilised in the distributed connectedess implementation side-by-side with the distributed infectiousness implementation. The vertical white and red dotted lines added to the gamma distribution plots mark mean values for the susceptible (blue) and infected (red) subpopulations, respectivelly.

The model with heterogeneity in infectiousness produces identical outputs to the homogeneous model for the same parameter values because infectiousness is not under selection by the force of infection. Heterogeneity in connectedness implies a positive correlation between the dispositions to acquire infection and transmit to others. Effectively, this results in infectiousness being selected indirectly via acquisition of infection, enhancing the effects observed under heterogeneous susceptibility alone.

It is evident from this exercise that knowing the extent of variation in susceptibility present in a population is essential if models are to be predictive. Variation in infectiousness does not affect predictability unless it is correlated with susceptibility, such as in the case of heterogeneity in connectedness. Susceptibility involves a probability of response to a stimulus (i.e. become infected given a pathogen challenge) and therefore cannot be measured directly. This obstacle, which may be part of the reason behind the widespread adoption of homogeneous models, is starting to be overcome by specific study designs that recognise the need for unpacking exposure gradients [7, 20, 12, 5, 17, 13, 8] as explicit experimental or observational dimensions.

### Co-circulation of multiple pathogen lineages

Many pathogens appear structured into multiple genetic lineages which are simultaneously maintained within host populations. Mathematical models, typically tied to lineages being homogeneous static entities [14], have invoked interactions between strains for stabilising coexistence. Here I argue that unobserved host heterogeneity in susceptibility within lineages alone can stabilise coexistence of multiple lineages.

Consider a discretised version of the heterogeneous susceptibility *SI* model:

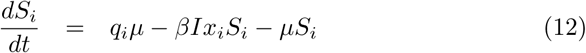

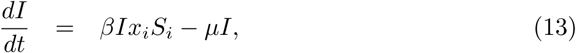

where *x*_*i*_, for *i* = 1, *…*, *n*, are the susceptibility factors of hosts *S*_*i*_ that enter the system as a fraction *q*_*i*_ of all births, purporting a distribution with mean ⟨*x*⟩ = Σ_*i*_ *q*_*i*_*x*_*i*_ *=*1, variation ⟨(*x* − 1)^2^⟩= Σ_*i*_ *q*_*i*_(*x*_*i*_ − 1)^2^, and coefficient of variation 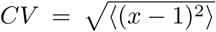, which will be treated as a varying parameter. The basic reproduction number is *R*_0_ = *β/µ*.

The extension of the model to *N* pathogen lineages, each characterised by an independent susceptibility distribution, is straightforward although the notation becomes cumbersome:

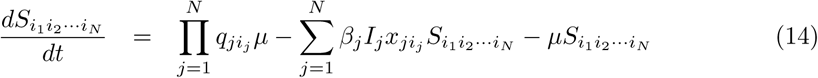

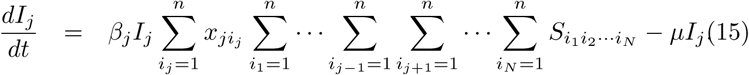

where *β*_*j*_, for *j* = 1, *…*, *N*, is the effective contact rate between hosts infected by species *j* and susceptible hosts, 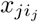, for *i*_*j*_ = 1, *…*, *n*_*j*_, are the susceptibility factors of hosts *S*_*…*_*i_j_*_*…*_, who enter the system as fractions *q*_*ji*_*j* of all births, purporting distributions with mean 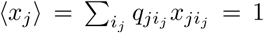, variance 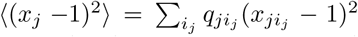, and coefficients of variation 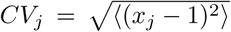 treated as varying parameters. The lineage-specific basic reproduction numbers are 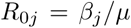. This system accommodates an *N*-lineage coexistence region with all 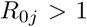. This region has a simple geometry in the 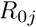 space. In the special case where the host population is homogeneously susceptible to lineage 1, it is bounded by the hyperplanes 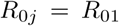, for *j* = 2, *…*, *N*, and by a hypersurface that can be obtained by setting to zero the abundance of 1 (shown in Figure 3 for 2 and 3 lineages). This coexistence region persists when we allow for heterogeneous susceptibility to lineage 1 as well.

**Figure 3:**
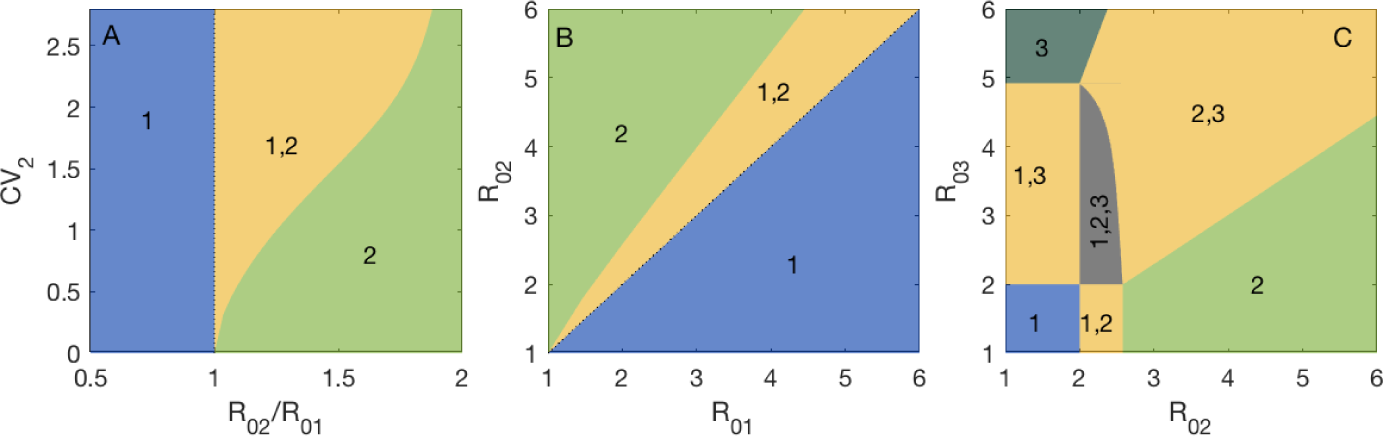
Stable coexistence of microbial lineages colonising a host population. Model (14)-(15) was solved analytically with two (A, B) and three (C) lineages. Yellow regions represent conditions for 2-lineage stable coexistence as indicated, while 3-lineage coexistence is found in the grey zone (C). Other parameters: (A, B, C) *CV*_1_ = 0; (B, C) *CV*_2_ = 1; (C) *CV*_3_ = 2 and *R*_01_ = 2.

## 5 Outlook

Selection within cohorts is ubiquitous in living systems, with manifold manifestations in any study that involves counting the individuals that constitute a population over time, across environments or experimental conditions. Whether we refer to populations of animals, microbes, or cells, the idea that in every observational or experimental study there is always a degree of unobserved heterogeneity that can reverse the direction of our conclusions is unsettling, but the issue can be tackled by general mathematical formalisms that account for it [1, 4, 10, 11, 16, 17, 22, 23] combined with study designs that enable its estimation [7, 8, 9, 12, 13, 20]. Collectively, the phenomenon appears to explain a wide range of reported discrepancies between studies and contribute to resolve decade-long debates, such as why vaccines appear less efficacious where disease burdens are high [5] and whether niche mechanisms need to be invoked to explain the levels of biodiversity observed in nature [6].

In addition to its omnipresence in studies that deal with populations explicitly, selection within cohorts may also play a central part in much debated issues surrounding accuracy and reproducibility of biological results more generally. Among aspects of research reproducibility discussed in the literature, those that pertain to methodology have focussed on how sample sizes must be sufficiently large to ensure a satisfactory level of certainty on the conclusions [2] and how shuffling is necessary to randomise conditions [18]. Additional problems, however, may result from overlooking forms of selection that may be occurring throughout the experiment. Any count of responses to a stimulus is affected by selection bias (unless all individuals, or cells, or other units, in the experiment were perfectly identical, which they are not). This is particularly problematic when different treatments are setup with the intent of comparing how differently individuals respond in one experimental condition vs another. Since the levels of selection bias will generally differ between treatments, comparisons of direct counts are not accurate representations of how differently individuals respond. Similar arguments apply to comparisons between different runs of entire experiments and compromise reproducibility. This is a problem of accuracy which cannot be resolved by increasing sample size or randomisation, but rather by adding dimensions to the experimental design [7]. This methodological issue can induce not only quantitative deviations in experimental results, but also invert the conclusions altogether [12].

This paper conveys how a wide variety of phenomena can be alternatively described by population thinking (individuals are different and selection operates) or individual thinking (all individuals are average and additional processes must be invoked) reaching contrasting conclusions. It is thus imperative to understand which frame is most appropriate is each case, and this implies understanding which individual characteristics may be subject to selection and how to obtain realistic descriptions of their variability in a population. The concerns presented here are pervasive across life and social sciences and can only be tackled in tight alliance with the mathematical sciences.

## Acknowledgments

This research was supported by Fundação para a Ciência e a Tecnologia (IF/ 01346/2014). An earlier version of this paper was published in *CIM Bulletin* in commemoration of the Year of Mathematical Biology 2018.

